# Biomarker de-Mendelization: principles, potentials and limitations of a strategy to improve biomarker prediction by reducing the component of variance explained by genotype

**DOI:** 10.1101/428276

**Authors:** Alisa D Kjaergaard, Stig E Bojesen, Børge G Nordestgaard, Julia S Johansen, George Davey Smith

**Affiliations:** Department of Clinical Biochemistry, Aarhus University Hospital, Denmark; Department of Clinical Biochemistry, Herlev and Gentofte Hospital, Copenhagen University Hospital, Denmark; The Copenhagen City Heart Study; Bispebjerg and Frederiksberg Hospital, Copenhagen University Hospital, Denmark; The Copenhagen General Population Study, Herlev and Gentofte Hospital, Copenhagen University Hospital, Denmark; Faculty of Health and Medical Sciences, University of Copenhagen, Denmark; Departments of Oncology and Medicine, Herlev and Gentofte Hospital, Copenhagen University Hospital, Denmark; MRC Integrative Epidemiology Unit, University of Bristol, United Kingdom

**Author notes:** Correspondence to: Alisa D Kjaergaard.

**Keywords:** YKL-40, CHI3L1, rs4950928, lung cancer, alcoholic liver disease, biomarker de-Mendelization

## Abstract

In observational studies, the Mendelian randomization approach can be used to circumvent confounding, bias and reverse causation, and to assess a potential causal association between a biomarker and risk of disease. If, on the other hand, a substantial component of variance of a non-causal biomarker is explained by genotype, then genotype could potentially attenuate the observational association and the strength of the prediction. In order to reduce the component of variance explained by genotype, an approach that can be seen as the inverse of Mendelian randomization - biomarker de-Mendelization - appears plausible.Plasma YKL-40 is a good candidate for demonstrating principles of biomarker de-Mendelization because it is a non-causal biomarker with a substantial component of variance explained by genotype. This approach is an attempt to improve the observational association and the strength of a predictive biomarker; it is explicitly not aimed at detection of causal effects.

We studied 21 161 individuals form the Danish general population with measurements of YKL-40 concentration and rs4950928 genotype. Four different methods for biomarker de-Mendelization are explored for alcoholic liver cirrhosis and lung cancer.

De-Mendelization methods only improved predictive ability slighly. We observed an interaction between genotype and markers of developing disease with respect to YKL-40 concentration.

Even when genotype explains 14% of the variance in a non-causal biomarker, we found no useful empirical improvement in risk prediction by biomarker de-Mendelization. This could reflect the predictive interaction between genotype and disease development being removed which counterbalanced any beneficial properties of the method in this situation.

## Introduction

Observational studies are prone to confounding, bias and reverse causation, which limits their ability to identify causal associations. Indeed, there are numerous examples of observational associations, that failed to be reproduced in randomized controlled trials [1,2].

In order to circumvent confounding and reverse causation, and to asses a potential causal association between a biomarker and risk of disease, the Mendelian randomization approach can be used. Genetic variants which are associated with biomarker concentration are employed to strengthen causal inference [3–8], and, when additional assumptions are satisfied, allow estimation of the magnitude of causal effects [9]. Mendelian randomization utilizes Mendel’s first and second law of inheritance stating that there is no segregation distortion and that a random assortment of alleles takes place at the time of conception [10]. Consequently, genotype is largely unconfounded, with confounders being distributed equally across genotype [11]. Reverse causation is impossible as genotype is established at conception and is constant throughout life. Causal effects are examined with randomization, which is by genotype in the Mendelian randomization approach, whereas it is by treatment and placebo in the controlled randomized trial. Generally, Mendelian randomization analyses produce clearer results when a substantial component of variance of a causal biomarker is explained by genotype. On the other hand, when a substantial component of variance in a non-causal biomarker is explained by genotype, then genotype could potentially attenuate the observational association and the strength of a predictive biomarker, because the component of variance accounted for by genotype will not predict disease. In order to reduce the influence of the component of variance explained by genotype, an approach that can be seen as the inverse of Mendelian randomization seems plausible. Reducing the quantitative contribution of genetic variants that influence the concentration of a non-causal biomarker - or “biomarker de-Mendelization” - could improve the utility of a biomarker intended to be predictive of disease occurrence, as in screening for disease or identifying a high risk group for preventive interventions. That prediction could in principle be improved through such an approach has been alluded to previously, [12, 13] and has been explored, with variable findings, in relation to genetic variants that influence prostate specific antigen levels, but not risk of prostate cancer [14–16]. The approach has not been formalized, or different analytical methods compared, however. It is important to emphasize that this “biomarker de-Mendelization” is an attempt to improve the observational association and the strength of a predictive biomarker. It is explicitly not aimed at detection of causal effects.

The optimal candidates for biomarker de-Mendelization are biomarkers with strong genotype-phenotype correlation, and where only phenotype (biomarker concentration) and not genotype is associated with risk of disease. Plasma YKL-40 is such a biomarker. A promoter SNP rs4950928 in the gene for YKL-40 *(CHI3L1)* explains 14% of the variation, and genotype is associated with doubling and tripling in plasma YKL-40 concentrations compared with the reference genotype [17–19]. Furthermore, this SNP is likely functional since MYC/MAX transcriptional factors bind better to the major than the minor allele of rs4950928 [20]. The better binding to the major allele results in increased *CHI3L1* transcription [21], and consequently higher plasma YKL-40 concentrations for each additional major allele [17–19]. Thus, this SNP only affects concentrations of plasma YKL-40, and not the protein function. This, and the high frequency of the rare allele makes this SNP a very good genetic indicator of lifelong increased plasma YKL-40 concentrations [18].

We have previously reported that a baseline increased plasma YKL-40 was associated with up to a 41-fold increased risk of alcoholic liver disease, and less than a 2-fold increased risk of lung cancer, with no evidence to support causal relationships [17, 19]. This suggests that plasma YKL-40 is a marker rather than a causal factor in the development of these diseases. We would therefore expect that reducing the component of variance explained by genotype would improve the observational association and the strength of this predictive biomarker.

We tested the hypothesis that biomarker de-Mendelization, compared to conventional analyses, improves observational associations and the strength of YKL-40 as a predicitive biomarker for alcoholic liver cirrhosis and lung cancer. We chose these two endpoints because risks of both were associated with plasma YKL-40 concentrations, but at very different magnitudes.

For this purpose, we performed cohort studies in 21 161 individuals from the Danish general population who had measurements of both plasma YKL-40 concentration and rs4950928 genotype. First, we examined association between YKL-40 concentration and risk of alcoholic liver cirrhosis and lung cancer. Next, we compared four different biomarker de-Mendelization methods to a conventional Cox regression analysis. Finally, we evaluated YKL-40 as a disease risk predictor in terms of area under the receiver operating curve with and without biomarker de-Mendelization.

## Materials and Methods

### Participants

We used two independent cohorts of the Danish general population: the 1991–1994 examination of the Copenhagen City Heart Study (CCHS) [22] and the Copenhagen General Population Study (CGPS) [17–19] initiated in 2003 and with ongoing enrollment. All participants were whites of Danish descent. All individuals only participated in one study.

Participants filled out a self-administered questionnaire which was completed together with an investigator on the day of attendance. A physical examination was performed, and blood was drawn for immediate biochemical analyses, as well as for storage at −80°C for later DNA and biochemical analyses.

### The CCHS Study

The CCHS is a prospective cohort study initiated in 1976, with emphasis on participants aged 20–100. The original cohort (14 223 participants) was invited to participate in 4 re-examinations (in 1981–83, 1991–94, 2001–2003 and 2011–2013) together with a supplemental cohort of younger individuals (<50 years old) at each examination. We only used the data from the 1991–1994 examination of the CCHS because DNA was collected at this point. Measurements of both plasma YKL-40 concentrations and *CHI3L1* rs4950928 genotype as well as information on diagnoses of alcoholic liver cirrhosis and lung cancer were available in 7981 individuals.

### The CGPS Study

The CGPS is a prospective cohort study initiated in 2003 with ongoing enrollment and emphasis on participants aged 20–100. Measurements of both plasma YKL-40 concentrations and *CHI3L1* rs4950928 genotype were available in 13 180 individuals, examined 2003 through 2011.

### Ethics

The studies were conducted according to the Declaration of Helsinki and approved by Herlev Hospital and a Danish scientific ethical committee (No. 100.2039/91 and 01–144/01). All participants gave written informed consent.

### Endpoints

Diagnoses of alcoholic liver cirrhosis and lung cancer were defined according to the World Health Organization International Classification of Diseases; the 8th and 10th edition for alcoholic liver cirrhosis (ICD8: 571.09; ICD10: K70.3) and the 7th and 10th edition for lung cancer (ICD7: 162–164; 462; 562; 962–964. ICD10: C33–34; C37–38). Diagnoses were obtained from the national Danish Patient Registry, the national Danish Causes of Death Registry, and the national Danish Cancer Registry, with a very high degree of completeness [19]. Date of death was obtained from the national Danish Civil Registration System. We did not lose track of any participant.

### YKL-40 analysis

Plasma concentrations of YKL-40 were determined in duplicates in samples frozen for 12–15 years at −80°C in the CCHS (N=7981) and in fresh samples (N=3205) or samples frozen for 7 years at −80°C (N=9975) in the CGPS by a commercial two-site, sandwich-type enzyme-linked immunosorbent assay (ELISA) (Quidel Corporation, San Diego, California). This assay uses streptavidin-coated microplate wells, a biotinylated-Fab monoclonal capture antibody, and an alkaline phosphatase-labeled polyclonal detection antibody. The recovery of the ELISA was 102% and the detection limit was 20 μg/L. The intra-assay coefficients of variation were 5% (at 40 μg/L) and 4% (at 104 μg/L and 155 μg/L). The inter-assay coefficient of variation was <6%. There is no circadian variation in plasma levels of YKL-40 [23], and it remains stable regardless of repetitive freezing and thawing [24].

### Genotype

Genotyping was performed using TaqMan assays (Applied Biosystems by Life Technologies Corporation, Carlsbad, CA, USA). Genotypes were in Hardy-Weinberg equilibrium (P=0.19, Table S1 in Appendix). We used a forward (AGT TCC CAT AAA AGG GCT GGT TT) and a reverse (CCC AGG CCC TGT ACT TCC TTT ATA T) primer for the PCR amplification and common (CTCCCCCACGCGGC) and variant (ACTCCCCGACGCGGC) probes to determine genotype.

### Covariates

Participants reported on present and past smoking habits and weekly alcohol consumption in the questionnaire. A pack-year was defined as 20 cigarettes smoked per day for 1 year (or equivalent). Alcohol consumption was defined as number of drinks per week, where 1 drink is equivalent to 12 g of alcohol. Body mass index in kg/m^2^ was calculated from measured weight in kg divided by measured height in meters squared. We used alanine aminotransferase, alkaline phosphatase, albumin, erythrocyte mean corpuscular volume, bilirubin, γ-glutamyl transpeptidase, coagulation factors II, VII, and X, pancreas amylase, C-reactive protein (CRP), and fibrinogen as markers of possible liver damage. These were measured by use of standard hospital assays subjected to quality control assessment [17]. A complete set of data on all covariates was not available for 1280 participants (6%), and these were imputed using multiple imputation; however, if only individuals with complete data were analysed, results were similar to those reported.

### Statistical analysis

We used statistical software STATA version 13. Comparison of groups was performed by χ^2^ and Cuzick’s non-parametric test for trend. No corrections for multiple comparisons were performed in these exploratory analyses.

Mean and median plasma YKL-40 concentration increases exponentially with increasing age (Figure 1). In all analyses, YKL-40 was therefore corrected for age. We calculated YKL-40 percentile categories (0–33%, 34–66%, 67–90%, 91–95% and 96–100%) within each 10-year age group (decade), and pooled the respective percentile categories together across age groups. We have previously used these percentile categories in order to evaluate both tertiles in the low range and extremely high plasma YKL-40 concentrations in smaller groups within the top tertile [17,19].

**Fig. 1.**
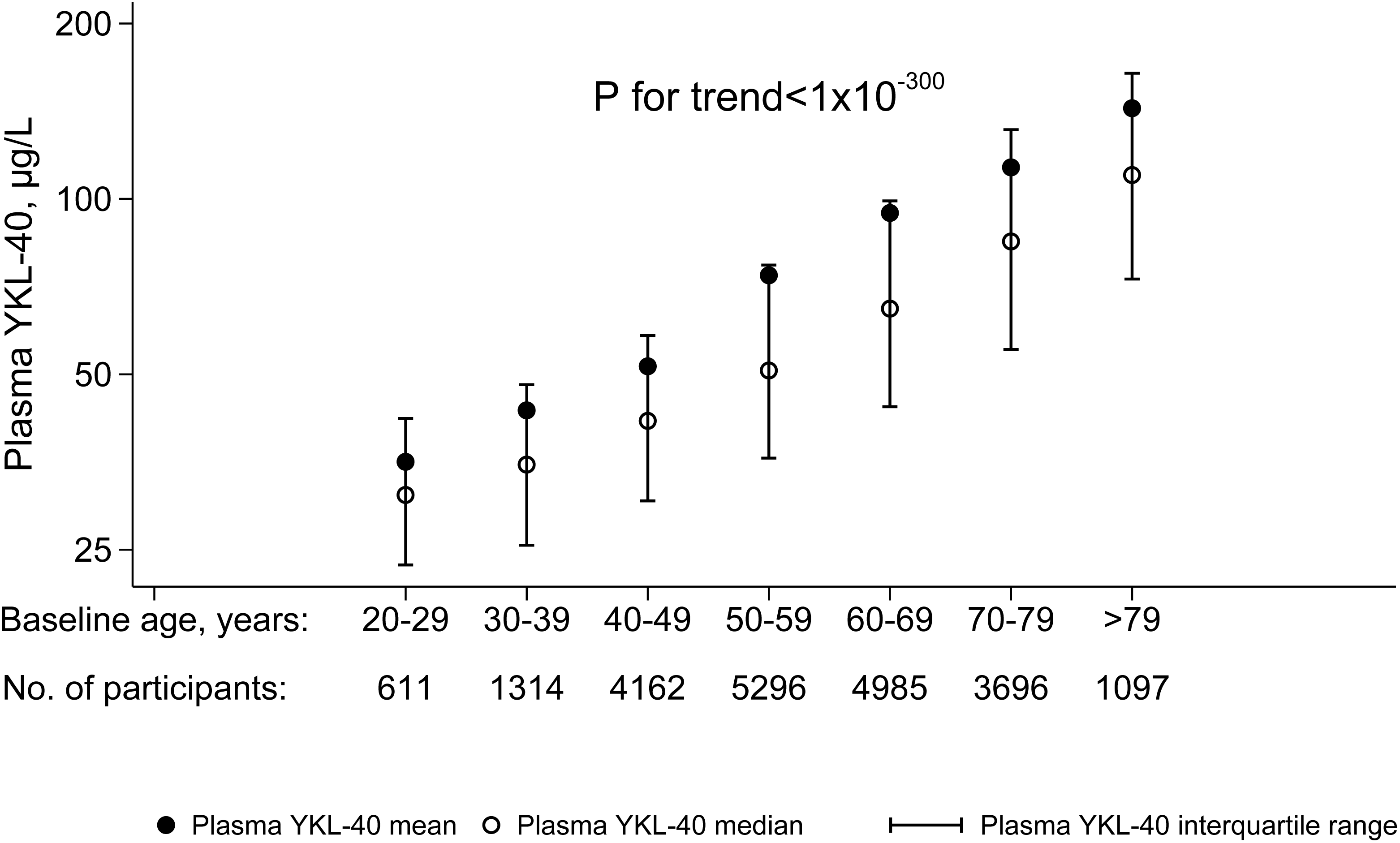
Plasma YKL-40 and age. Plasma YKL-40 concentration across increasing age categories in 10-year age groups. P-value is for Cuzick’s nonparametric test for trend.

Risk estimates and corresponding 95% confidence intervals for alcoholic liver cirrhosis and lung cancer were computed as hazard ratios by Cox regression analysis. We did not correct for regression dilution bias.

We found no marked violation of proportional hazards assumption graphically (by plotting –ln(survival probability) against ln(analysis time)) or by Schoenfeld residuals. In the Cox analysis, we used age as time scale. Participants entered at the time of blood sampling, and were followed until event, death or end of follow-up May 10^th^ 2011 for alcoholic liver cirrhosis and December 31^st^ 2011 for lung cancer, whichever came first. Emigrated participants (n=104) were censored at the time of emigration. Participants with events prior to blood sampling were excluded from the respective analyses.

### Biomarker de-Mendelization methods

We performed four different methods for biomarker de-Mendelization: genotype adjusted, meta-analysis, residual and genotype stratified.

For the genotype adjusted method, we used genotype (=number of C alleles) as a continuous adjustment variable in the Cox regression analysis. For the meta-analysis method, we performed meta analyses on estimates derived from the analyses within each genotype separately. For the residual method, we corrected plasma YKL-40 concentrations for genotype, by using the residuals from linear regression of YKL-40 on rs4950928 (Figure 2). For the genotype stratified method, we calculated YKL-40 percentile categories for each genotype separately, and pooled the respective categories together.

**Fig. 2.**
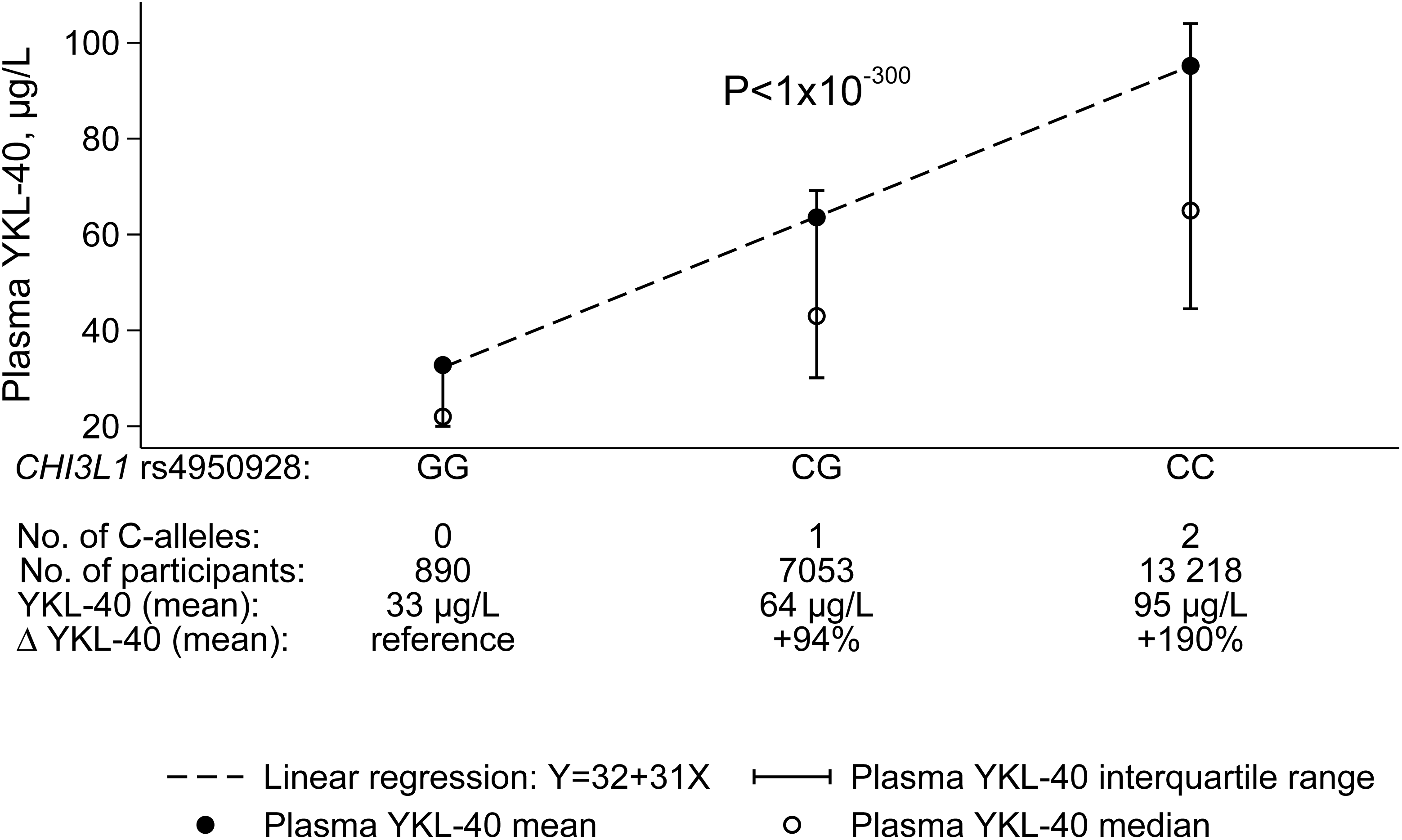
Genotype-phenotype association. Plasma YKL-40 concentration according to *CHI3L1* rs4950928 genotype. P-value is for Cuzick’s nonparametric test for trend.

In the multifactorially adjusted analyses, we further adjusted for sex, cumulative smoking (pack-years), alcohol consumption (number of drinks per week, 1 drink ≈ 12 g alcohol), body mass index (kg/m^2^) and study population (CCHS or CGPS) for both endpoints; as age was the underlying time-scale, age is automatically adjusted for.

Moreover, we evaluated YKL-40 as a risk predictor (area under the receiver operating curve) of disease with and without biomarker de-Mendelization in a logistic regression model.

Finally, we explored whether genotype (rs4950928) showed differing association with plasma YKL-40 according to markers of disease development. To calculate P for interaction, we used a likelihood ratio test after linear regression between the models with and without an interaction term between genotype and marker of disease development.

## Results

The combined study included 21161 individuals from the Danish general population with measured YKL-40 concentrations and rs4950928 genotype. From 1976 through 2011, with information on all participants, 96 individuals developed alcoholic liver cirrhosis, and 434 developed lung cancer (prevalent disease, Table S1 in Appendix). However, we only considered incident events (58 cases of alcoholic liver cirrhosis and 401 cases of lung cancer; Figures 3, 4 and 5, and Figures S1 and S2 in Appendix) for the Cox regression and ROC curves, as participants with events prior to blood sampling were excluded from these analyses, because we wanted to test the prospective predictive ability of YKL-40.

**Fig. 3.**
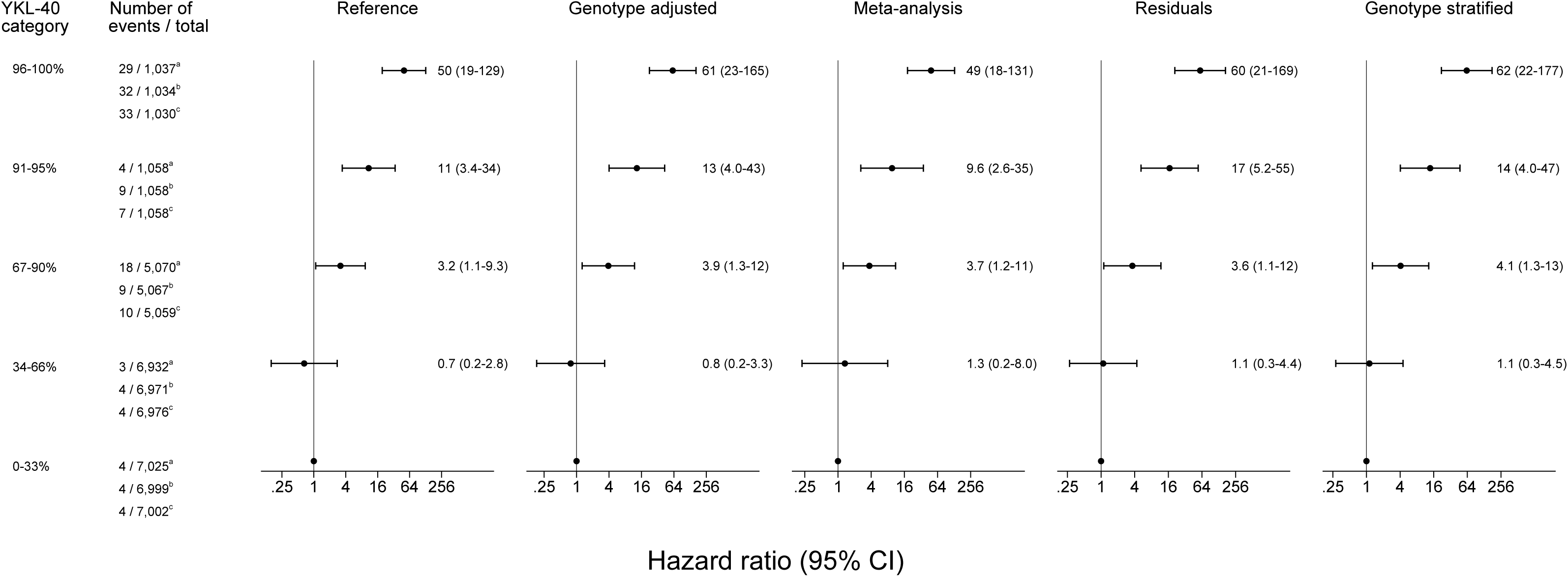
Age and sex adjusted hazard ratios for alcoholic liver cirrhosis according to plasma YKL-40 percentile category. YKL-40 percentile categories were calculated based on measured YKL-40 concentrations for the reference method as well as for genotype adjusted and meta-analysis methods. Because YKL-40 percentile categories were calculated based on the residuals for the residual method, and for each genotype separately for the genotype stratified method, the distribution of participants across YKL-40 percentile categories differed slightly between these three different ways of calculating YKL-40 percentile categories ^a^Reference, genotype adjusted and meta-analysis methods. ^b^Residuals method. ^c^Genotype stratified method.

**Fig. 4.**
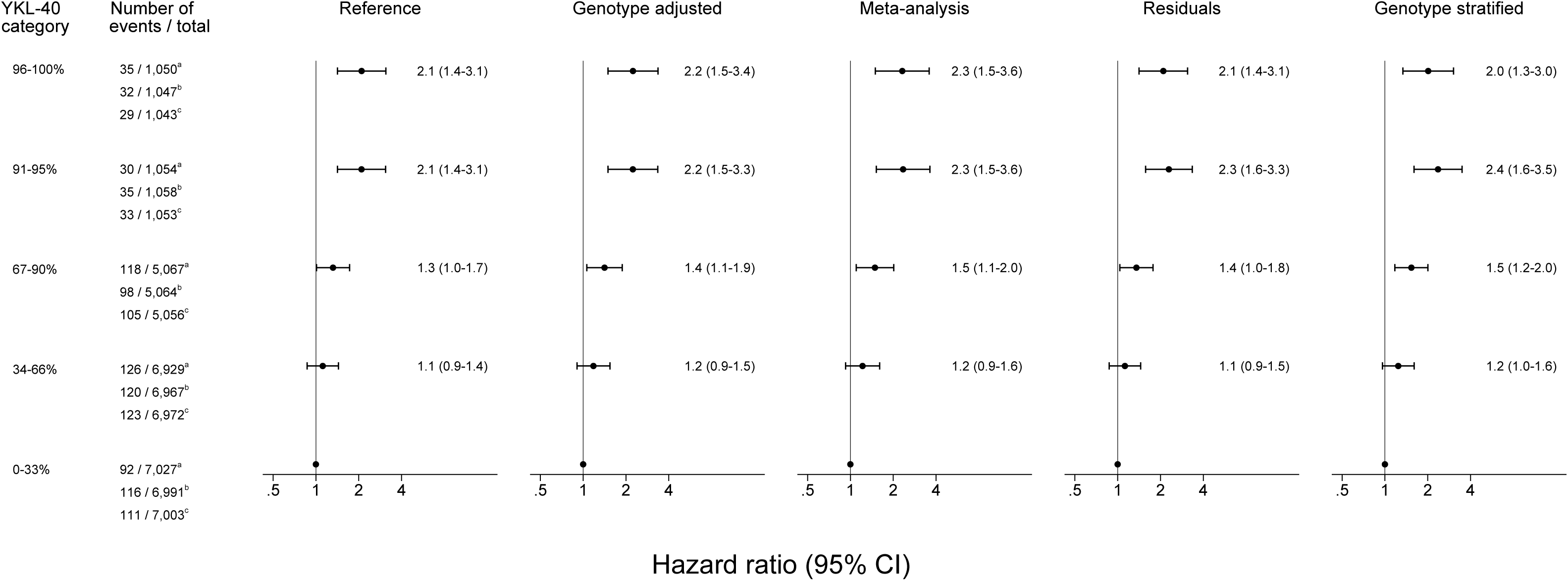
Age and sex adjusted hazard ratios for lung cancer according to plasma YKL-40 percentile category. YKL-40 percentile categories were calculated based on measured YKL-40 concentrations for the reference method as well as for genotype adjusted and meta-analysis methods. Because YKL-40 percentile categories were calculated based on the residuals for the residual method, and for each genotype separately for the genotype stratified method, the distribution of participants across YKL-40 percentile categories differed slightly between these three different ways of calculating YKL-40 percentile categories ^a^Reference, genotype adjusted and meta-analysis methods. ^b^Residuals method. ^c^Genotype stratified method.

**Fig. 5.**
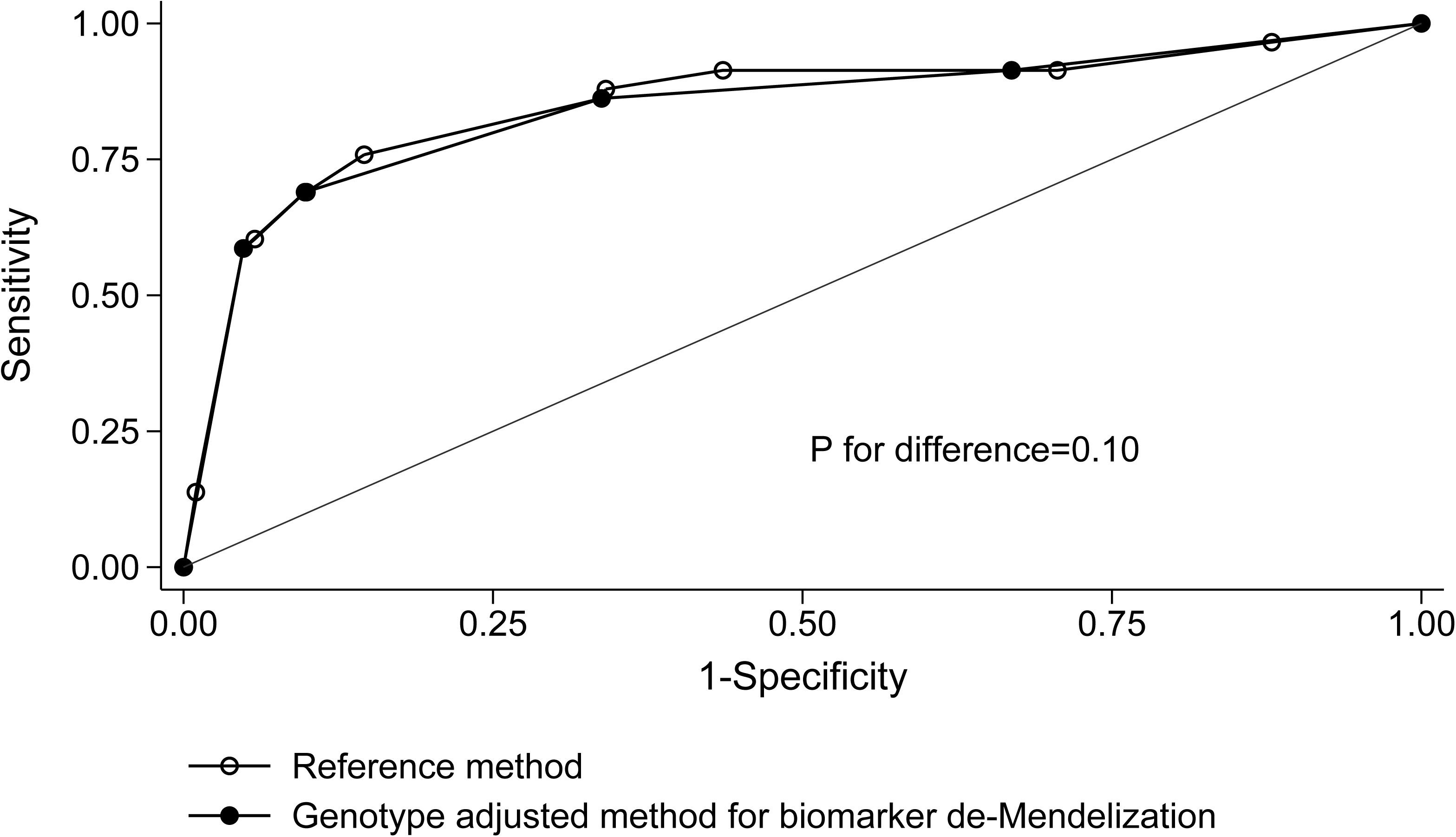
Receiver operating characteristics (ROC) curves for alcholic liver cirrhosis with and without biomarker de-Mendelization. P-value is for χ^2^ from STATA’s roccomp command, testing whether the areas under the curves are equal.

*CHI3L1* rs4950928 genotypes were in Hardy-Weinberg equilibrium (P=0.19, Table S2 in Appendix), with a C allele frequency of 79%. Baseline characteristics as a function of YKL-40 percentile categories, alcoholic liver disease, lung cancer and rs4950928 genotype are shown in Table 1 and Tabels S1 and S2 in Appendix; YKL-40 concentrations and later disease event were strongly associated with most covariates, while genotype was not.

**Table 1.** Characteristics of participants. Baseline values collected at the 1991–1994 examination of the Copenhagen City Heart Study, and the 2003–2011 examination of the Copenhagen General Population Study, and expressed as numbers of participants, frequencies, or medians (interquartile ranges). NA=not applicable. P-values are from Cuzick’s non-parametric test for trend. ^a^Plasma YKL-40 concentrations in percentile categories overlap for some categories due to correction for age.

|  | Plasma YKL-40 percentile category |  |  |  |  | P-value |
| --- | --- | --- | --- | --- | --- | --- |
|  | 0-33% | 34-66% | 67-90% | 91-95% | 96-100% |  |
| Number of participants, N | 6994 | 6989 | 5067 | 1059 | 1052 | NA |
| Women, % | 58 | 56 | 54 | 54 | 41 | $2 \times 10^{-21}$ |
| Age, years | 58 (47-68) | 58 (48-68) | 58 (48-69) | 58 (48-69) | 58 (48-68) | 0.01 |
| Alcohol consumption, weekly drinks | 6 (2-12) | 7 (3-14) | 9 (3-17) | 11 (4-20) | 16 (7-30) | $2 \times 10^{-178}$ |
| Cumulative smoking, pack-years | 7 (0-25) | 9 (0-28) | 11 (0-31) | 16 (0-35) | 23 (2-41) | $3 \times 10^{-65}$ |
| Body mass index, kg/m<sup>2</sup> | 25.0 (22.7-27.7) | 25.3 (22.9-28.2) | 25.7 (23.0-28.6) | 26.0 (23.4-28.9) | 25.9 (23.1-29.4) | $2 \times 10^{-23}$ |
| Plasma YKL-40, µg/L<sup>a</sup> | 32 (25-40) | 55 (43-73) | 91 (67-124) | 163 (108-207) | 267 (188-473) | NA |

### Plasma YKL-40 and age

Plasma YKL-40 concentrations increase exponentially with increasing age (Figure 1), but calculating YKL-40 percentile categories within each 10-year age group resulted in similar age distributions (median and interquartile range) across the five YKL-40 percentile categories: 0–33%, 34–66%, 67–90, 91–95% and 96–100% (Table 1).

### Genotype-phenotype association

Arithmetic mean for plasma YKL-40 was 33 μg/L for individuals with the GG genotype, 64 μg/L for the CG genotype (doubling from 33 μg/L), and 95 μg/L (tripling from 33 μg/L) for the CC genotype, corresponding to a linear regression of YKL-40 on rs4950928 genotype of (Figure 2.): Predicted YKL-40=32+31 x number of C-alleles (P<1×10^−300^, r^2^=0.14).

### Plasma YKL-40 and risk of disease: biomarker de-Mendelization

Higher percentile of plasma YKL-40 was associated with higher risk of alcoholic liver cirrhosis and lung cancer (first panels in Figures 3 and 4, Figures S1 and S2 in the Appendix).

Compared to the 0–33 percentile category, reference hazard ratios for alcoholic liver cirrhosis were 11 (95%CI:3.4–34) and 50 (19–129) for the 91–95 and 96–100 percentile categories, respectively. Corresponding hazard ratios for de-Mendelization methods were 13 (4.0–43) and 61 (23–165), 9.6 (2.6–35) and 49 (18–131), 17 (5.2–55) and 60 (21–169) and, 14 (4.0–47) and 62 (22–177) for the genotype adjusted method, meta-analysis, residuals and genotype stratified method, respectively (Figure 3). Multifactorially adjusted analyses showed similar results (Figure S1 in Appendix).

Compared to the 0–33 percentile category, reference hazard ratios for lung cancer were 2.1 (95%CI: 1.4–3.1) and 2.1 (1.4–3.1) for 91–95 and 96–100 percentile category, respectively. Corresponding hazard ratios for de-Mendelization methods were 2.2 (1.5–3.3) and 2.2 (1.5–3.4), 2.3 (1.5–3.6) and 2.3 (1.5–3.6), 2.3 (1.6–3.3) and 2.1 (1.4–3.1), and 2.4 (1.6–3.5) and 2.0 (1.3–3.0) for the genotype adjusted method, meta-analysis, residuals and genotype stratified method, respectively (Figure 4). Multifactorially adjusted analyses showed similar results (Figures S1 and S2 in the Appendix).

### Plasma YKL-40 as a predictor of disease: biomarker de-Mendelization

As there were no substantial differences between the de-Mendelization methods, we performed analyses only for the genotype adjusted method. The areas under the reciever operating characteristics curves (AUC-ROC) hardly differed between the reference and de-Mendelization method (AUC-ROC: 0.84 and 0.85, P for difference =0.10; Figure 5) for alcoholic liver cirrhosis. Multifactorially adjusted analyses showed similar results (data not shown). For lung cancer, the AUC-ROCs were lower and similar for the reference and de-Mendelization methods (data not shown).

Compared to G-allele, the C-allele (YKL-40 increasing allele) was associated with an even greater increase in plasma YKL-40 concentration in participants who developed disease after blood sampling, and in participants who had highest tertiles of risk factors and potential markers of disease (Figures S3-S5 in Appendix). This was most pronounced for alcoholic liver cirrhosis (P for interaction=4×10^−11^), γ-glutamyl transpeptidase (P=4×10^−11^), C-reactive protein (P=6×10^−6^), and alcohol consumption (P=5×10^−4^) (Figures S3-S5 in Appendix).

## Discussion

In this demonstration of the principles of biomarker de-Mendelization we compared four different de-Mendelization methods to a usual Cox regression analysis. Studying 21161 individuals from the general population, we found no useful empirical improvement in risk prediction by biomarker de-Mendelization.

This was despite the fact that *CHI3L1* rs4950928 genotype explained 14% of the variance in plasma YKL-40 concentrations. Because most genetic instruments used in the Mendelian randomization studies explain far less than 14% of the variation in biomarker concentrations, it could be considered that it is unlikely that other studies would benefit from the proposed de-Mendelization methods. However, it is important to consider the assumptions that the approach depends upon. In the context of this study an important one is that genotype should not contribute to prediction through increasing the biomarker level in the presence signs of developing disease. In this study, we used a range of measures to detect this. If such an interaction exists, then this will improve the predictive ability of the biomarker, and removing the influence of genotype would abrogate this. Precisely such an interaction was evident. It seems that in this example, biomarker de-Mendelization had two opposing effects on risk prediction, as the component of variance accounted for by genotype could potentially both improve as well as abrogate risk prediction. This might explain our essentially null findings. However, this should be interpreted cautiously due to few events among participants with the rare GG genotype.

A second assumption is that the per-allele genotypic effect on the biomarker is reasonably linear. If not, then all but the stratification by genotype and meta-analysis approach would be sub-optimal. In our example, the genotypic effect is close to per-allele, and the simple adjustment for genotype as a continuous variable seems reasonable. An advantage of this method is its simplicity and intuitiveness compared to the other proposed de-Mendelization methods. Notably, no method was consistently substantially better than the others. Interestingly, de-Mendelization by meta-analysis method showed no improvement in risk prediction for alcoholic liver cirrhosis, but the largest improvement in risk prediction for lung cancer. An explanation for this could be that participants with the GG genotype (4.2% of the population) were excluded from the meta-analyses for both alcoholic liver cirrhosis and lung cancer due to lack of events; i.e. the meta-analyses were performed on participants with the CG and CC genotypes only. This problem would be emphasized in studies where minor allele frequency is even smaller. However, this potential problem could be circumvented by increasing sample size.

Because YKL-40 increases exponentially with increasing age, plasma YKL-40 percentiles were corrected for age by 10-year age bands, but even though median and interquartile range of age were roughly the same across YKL-40 percentile categories, there was a slight increase in mean age (57.4, 57.8, 58.1, 58.2 and 58.1 years, P=0.01) with increasing YKL-40 percentile category. However, a finer age adjustment was not possible due to our sample size (N=21161) and because we wanted the methods to be comparable (same age adjustment), and some of the methods (meta-analysis and genotype stratified method) required a further subdivision with very few events in some of the groups.

In the residual method of de-Mendelization, we used the residuals after linear regression of YKL-40 on rs4950928; i.e. we subtracted 32, 63 and 94 μg/L from the measured YKL-40 concentrations for individuals with GG, CG and CC genotypes, respectively. This was done for all participants regardless of age, and the residuals are probably too small for the youngest and too large for the oldest of participants. This might have influenced our risk estimates. Thus, it seems that the genotype stratified method of de-Mendelization could be a more appropriate choice for correcting for genotype in our study. It would be interesting to see how these four de-Mendelization methods perform for a biomarker with a weaker age association.

Potential limitations of this study are all the limitations of the basic observational data and the interpretation of these apply to the biomarker de-Mendelization analyses (17–19].

In conclusion, in this prospective study of individuals from the Danish general population, we found no useful empirical improvement in risk prediction of alcoholic liver cirrhosis and lung cancer by YKL-40 biomarker de-Mendelization. This could reflect the predictive interaction between genotype and disease development being removed, which counterbalanced any beneficial properties of the method in this situation.

## Funding

This work was supported by University of Copenhagen and by grants from Chief Physician Johan Boserup and Lise Boserups Foundation, the Danish Heart Foundation, the Danish Medical Research Council, the Research Council at Herlev and Gentofte Hospital, Toyota-Fonden Denmark and Vera and Carl Johan Michaelsen’s Foundation. Quidel provided the study with some of the YKL-40 ELISA kits. The funders had no role in study design, data collection and analysis, decision to publish, or preparation of the manuscript. All authors declare no conflict of interest.

## Conflict of interest

The authors declare that they have no conflict of interest.

## Acknowledgements

Excellent technical assistance has been provided by Annette Lundgren-Beck, Anja Jochumsen, Birgit Hertz, Debbie Nadelmann, Dorthe Mogensen, Hanne Damm, Marianne Sørensen, Nina Dahl Kjersgaard, Rikke Mirsbach Henkel and Ulla Kjærullf-Hansen, all from Herlev Hospital. We thank the participants of the Copenhagen City Heart Study and the Copenhagen General Population Study for their willingness to participate.

